# Role of AMPK in Atrial Metabolic Homeostasis and Substrate Preference

**DOI:** 10.1101/2024.08.29.608789

**Authors:** Zeren Toksoy, Yina Ma, Leigh Goedeke, Wenxue Li, Xiaoyue Hu, Xiaohong Wu, Marine Cacheux, Yansheng Liu, Fadi G. Akar, Gerald I. Shulman, Lawrence H. Young

## Abstract

Atrial fibrillation is the most common clinical arrhythmia and may be due in part to metabolic stress. Atrial specific deletion of the master metabolic sensor, AMP-activated protein kinase (AMPK), induces atrial remodeling culminating in atrial fibrillation in mice, implicating AMPK signaling in the maintenance of atrial electrical and structural homeostasis. However, atrial substrate preference for mitochondrial oxidation and the role of AMPK in regulating atrial metabolism are unknown. Here, using LC-MS/MS methodology combined with infusions of [^13^C_6_]glucose and [^13^C_4_]β-hydroxybutyrate in conscious mice, we demonstrate that conditional deletion of atrial AMPK catalytic subunits shifts mitochondrial atrial metabolism away from fatty acid oxidation and towards pyruvate oxidation. LC-MS/MS-based quantification of acyl-CoAs demonstrated decreased atrial tissue content of long-chain fatty acyl-CoAs. Proteomic analysis revealed a broad downregulation of proteins responsible for fatty acid uptake (LPL, CD36, FABP3), acylation and oxidation. Atrial AMPK deletion reduced expression of atrial PGC1-α and downstream PGC1-α/PPARα/RXR regulated gene transcripts. In contrast, atrial [^14^C]2-deoxyglucose uptake and GLUT1 expression increased with fasting in mice with AMPK deletion, while the expression of glycolytic enzymes exhibited heterogenous changes. Thus, these results highlight the crucial homeostatic role of AMPK in the atrium, with loss of atrial AMPK leading to downregulation of the PGC1-α/PPARα pathway and broad metabolic reprogramming with a loss of fatty acid oxidation, which may contribute to atrial remodeling and arrhythmia.

## Introduction

Atrial fibrillation (AF) is the most common arrhythmia affecting more than 6 million people in the United States alone ^1^. AF is a well-described complication in patients with hypertension, valvular disease, and heart failure. Atrial structural remodeling, in the form of chamber dilation and interstitial fibrosis, contributes to conduction delays that predispose to the electrical re-entry that is important in the genesis of this arrhythmia ^2^. Furthermore, alterations in ion channels and gap junction proteins underlie the electrophysiological substrate and triggers that lead to the initiation and perpetuation of AF ^2^.

A metabolic basis of clinical AF has been suggested based on its increased incidence in patients with aging, diabetes mellitus, and obesity ^1^. Manifestations of atrial metabolic stress have been documented in both patients and experimental models of AF, including oxidative stress and mitochondrial dysfunction ^3,4^. Atrial tissue from patients with persistent AF have also demonstrated a shift from fatty acid to glucose metabolism transcripts, with increased triglyceride and beta-hydroxybutyrate content, decreased glucose transporter expression, and alterations in both glycolytic and fatty acid metabolism enzymes ^5-10^. A recent study showed increased glucose uptake in patients with persistent AF using positron emission tomography (PET) with 18-fluorodeoxyglucose imaging, that normalized after ablation reestablished sinus rhythm ^11^. A sheep model of sustained AF also suggested a shift from fatty acid to glucose oxidation based on analysis of metabolic enzymes and intermediates ^12^. These prior studies of AF have not evaluated physiologic substrate oxidation or preference for mitochondrial metabolism. Furthermore, the role of metabolic stress as an underlying root cause of AF is poorly understood.

A potential regulator of atrial metabolism in the heart is the AMP-activated protein kinase (AMPK). AMPK is a serine-threonine kinase that is highly sensitive to the cellular energy status, due to its rapid activation in response to ATP breakdown, which increases the AMP/ATP and ADP/ATP ratios ^13^. AMPK maintains energy homeostasis by regulating lipid and glucose metabolism, redox state, and protein synthesis ^14^. AMPK is expressed abundantly in the ventricular myocardium of the heart and has emerged as a potential target for the treatment of ischemia-reperfusion and heart failure ^14-18^. Although loss of AMPK activity significantly impairs the metabolic response to myocardial ischemia, it does not alter baseline cardiac metabolism in the ventricle ^17^.

The role of AMPK in the cardiac atria is not well understood. In atrial tissue from patients with intermittent or “paroxysmal” AF, there is an increase in phosphorylation of the Thr^172^ activation site in the α catalytic subunit of AMPK ^19^. Conversely, AMPK phosphorylation is decreased in patients with long-standing or “persistent” AF, although the mechanism for the blunted activation is unknown ^20^. Experimental studies, including our own, have shown that cardiac deletion of LKB1 (the upstream kinase for AMPK and other AMPK-related kinases) causes AF in mice, albeit in the context of left ventricular dysfunction and heart failure ^20,21^. More direct insight into the role of AMPK in the atria came from our recent investigation showing that conditional deletion of AMPK in the atria causes spontaneous AF with reprogramming of ion channel and connexin expression that was associated with both electrical and structural atrial remodeling ^22^.

Due to the potentially important homeostatic role of AMPK in preventing atrial remodeling and AF, the current work aimed to define the effects of AMPK deletion on atrial metabolism. We utilized a mouse model of atrial deletion that avoids the confounding effects of loss of AMPK in the heart ventricles ^22^, with the goal to study the autonomous role of AMPK in regulating atrial metabolism. Our results show that mice with atrial AMPK deletion have substantial metabolic reprogramming, characterized by defective fatty acid uptake and oxidation and a greater reliance on myocardial glucose utilization, reminiscent of left ventricular metabolism in the fetal heart and the metabolic shift observed in heart failure ^23,24^. These findings implicate AMPK as having a major homeostatic role in atrial metabolism.

## Results

### Atrial AMPK depletion causes atrial electrical remodeling in AMPK dKO mice

Mice with conditional atrial deletion of AMPK α1 and α2 catalytic subunits were generated using the sarcolipin promoter to drive Cre expression in the atria while avoiding ventricular AMPK deletion, as previously described ^22^. These atrial AMPK double (α subunit) knock out (AMPK dKO) mice were studied, together with littermate floxed AMPK controls not expressing Cre-recombinase, at 4-6 weeks of age. We confirmed that the atrial AMPK dKO mice exhibited over 90% reduction in AMPK α subunit expression in both left and right atria (Figure 1A). Moreover, the downstream phosphorylation of acetyl-CoA carboxylase (ACC) on its canonical AMPK target site Ser^79^, and raptor on its AMPK target site Ser^792^, were both significantly reduced in AMPK dKO mice compared to littermate control mice (Figure 1A). The highly selective nature of the atrial deletion in the heart was evident, with no change observed in ventricular AMPK α subunit expression or downstream pathway phosphorylation. The extent of AMPK pathway inactivation was comparable between the left and right atria. Nonetheless, we designed the studies to separately analyze metabolic alterations in the two atria to determine whether they had a differential metabolic response to loss of AMPK activity.

**Figure 1:**
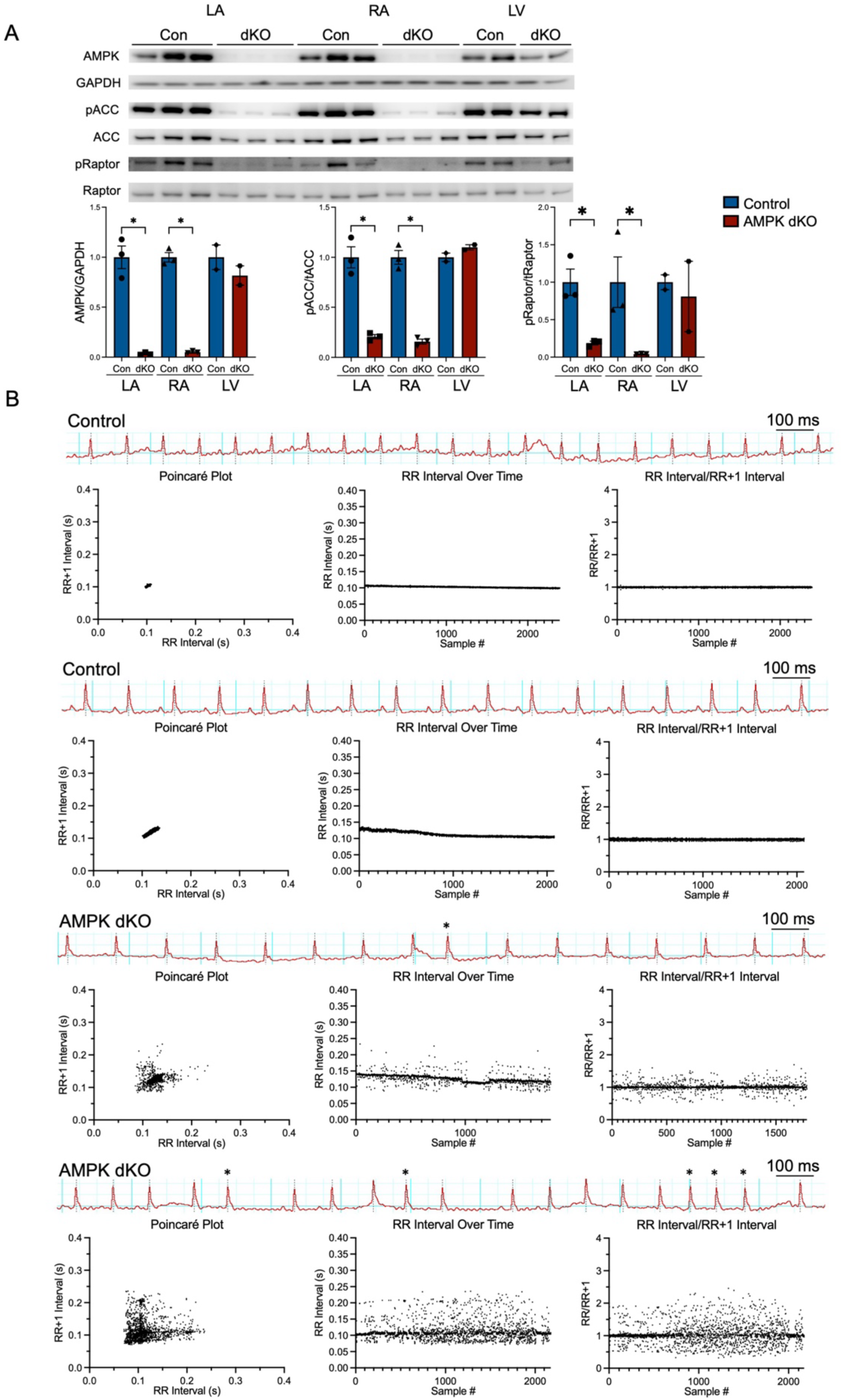
Atrial AMPK pathway inactivation and electrical remodeling in AMPK dKO mice. (A) Representative immunoblots showing the atrial content of AMPK, phospho-ACC Ser^79^, total ACC, phospho-raptor Ser^792^, and total raptor in homogenates from the left atria (LA), right atria (RA), and left ventricles (LV) of control (Con) and atrial AMPK α_1_α_2_ double knockout (AMPK dKO, also designated as dKO on axis labels) mice (n=3 per group). Quantification of AMPK is relative to GAPDH and quantification of phospho-proteins is relative to its respective total protein. (B) Representative electrocardiograms (ECGs) of control and AMPK dKO mice at 6 weeks of age with corresponding Poincaré, RR interval over time, and RR interval/RR+1 interval over time plots. Atrial ectopic complexes are noted by * symbols in the ECG tracings of AMPK dKO mice. Data are shown as mean ± SEM; Significance determined by unpaired 2-tailed Student’s *t* test. *p < 0.05 AMPK dKO vs. control

AMPK dKO mice exhibit sinus rhythm at 1 week of age, but start to display atrial ectopy starting at 2 weeks of age and intermittent spontaneous atrial fibrillation as early as 6 weeks of age ^22^. At 4-6 weeks, abnormal beat-beat heart rate variability was evident in AMPK dKO mice, in contrast to the minor physiological variability observed in control mice (Figure 1B). The greater heart rate variability was evident on Poincare plots, which correlate sequential RR intervals (RR vs. RR+1), as well as on plots of the RR intervals or the RR/RR+1 ratios over time (Figure 1B). The heart rate-variability was due to variation in the atrial rhythm, with the ECG tracings showing premature atrial complexes and short runs of atrial tachycardia (Figure 1B). We did not observe sustained atrial arrhythmia or atrial fibrillation at this early time point. Thus, we elected to do metabolic studies on mice at 4-6 weeks of age, which avoided the potential confounding effects of subsequent more persistent atrial arrhythmia on atrial metabolic remodeling.

### AMPK deletion alters atrial substrate oxidative metabolism

AMPK plays a critical role in regulating tissue metabolism, but its specific function in atrial metabolic homeostasis is unknown. Heart muscle generates ATP primarily through mitochondrial oxidative metabolism, with predominant oxidation of fatty acids under most conditions ^25^. Thus, we investigated the utilization of substrates for oxidative metabolism *in vivo* based on their incorporation into the tricarboxylic acid (TCA) cycle in the atrial AMPK dKO mouse model. The incorporation of glucose, ketones, and fatty acids into the myocardial TCA cycle was assessed during continuous intravenous infusions of ^13^C-labeled substrates in awake mice (Figure 2, A and B), using LC-MS/MS methods initially developed to study *in vivo* skeletal muscle metabolism ^26,27^.

**Figure 2:**
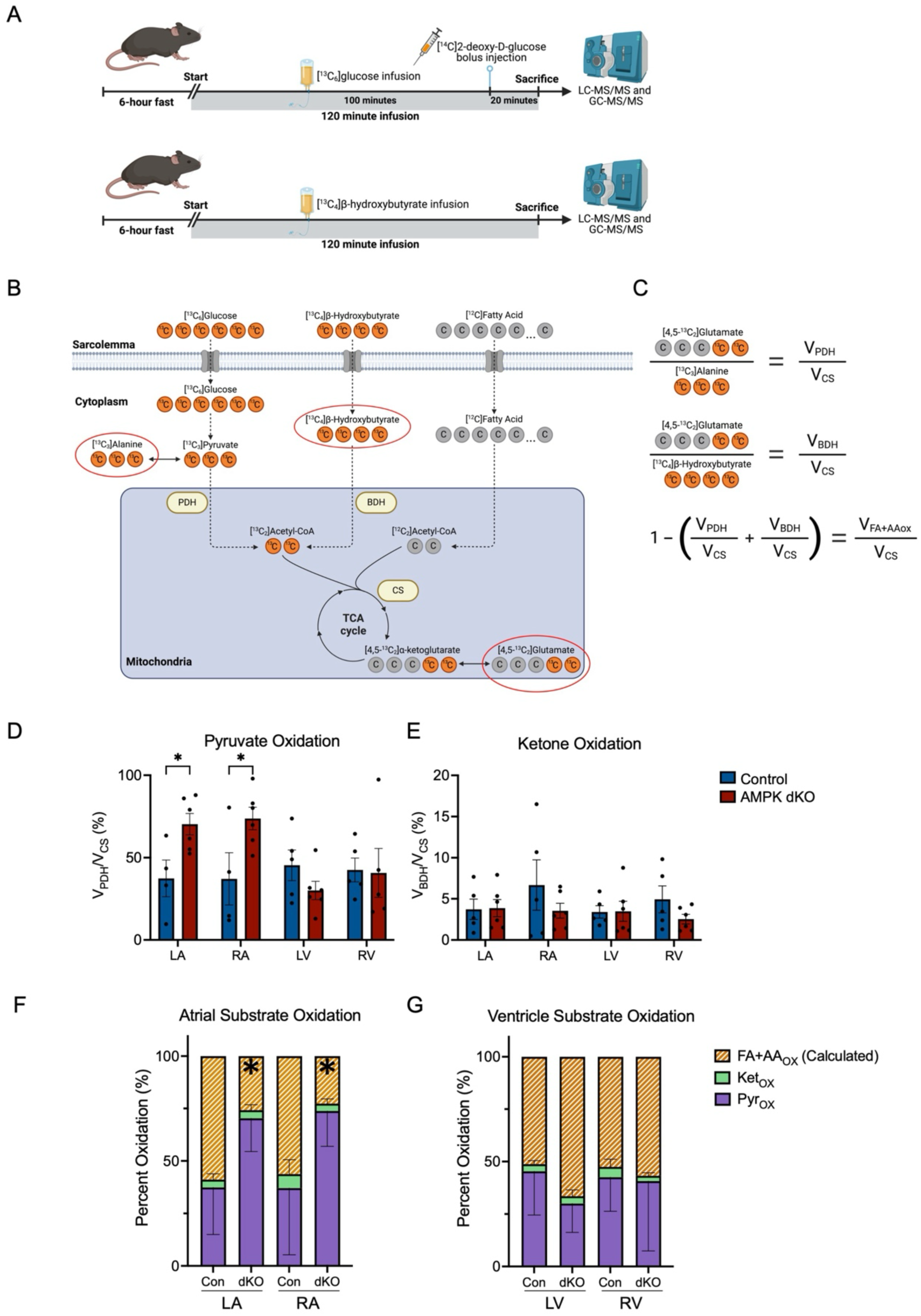
AMPK deletion alters atrial oxidative metabolism. (A) Schematic representation of metabolic flux studies. Following a 6-hour fast, control and atrial AMPK dKO conscious mice, at 4-6 weeks of age, were infused intravenously with either [^13^C_6_]glucose to assess myocardial pyruvate oxidation, or [^13^C_4_]β-hydroxybutyrate to assess myocardial ketone oxidation (n=4-6 per group). Glucose uptake was assessed after a bolus injection of [^14^C]2-deoxyglucose during the last 20 minutes of the [^13^C_6_]glucose infusions. (B) Schematic representation of isotopic labeling of atrial and ventricular metabolites during [^13^C] infusions: [^13^C]-labeled atoms denoted in orange and unlabeled atoms in grey. The [^13^C] enrichments of metabolites circled in red were measured by LC and GC-MS/MS. (C) Calculations used to assess the contributions of substrate entering the TCA cycle via either pyruvate dehydrogenase (PDH) during [^13^C_6_]glucose infusion (V_PDH_/V_CS_), via BDH during [^13^C_4_]β-hydroxybutyrate infusion (V_BDH_/V_CS_), or from fatty acid and amino acids (V_FA+AA_/ V_CS_). The relative rates are calculated as a ratio to the total flux into the TCA cycle via citate synthase (CS). (D and E) Pyruvate oxidation (V_PDH_/V_CS_) and ketone oxidation (V_BDH_/V_CS_) in the left atria (LA), right atria (RA), left ventricle (LV), and right ventricle (RV) of control and atrial AMPK dKO mice. (F and G) Relative contributions of pyruvate, ketone, and calculated fatty acid + amino acid oxidation expressed as a percent in each of the cardiac chambers. Data are shown as mean ± SEM; Significance determined by unpaired 2-tailed Student’s *t* test. *p < 0.05 AMPK dKO vs. control.

During intravenous infusions of [^13^C_6_]glucose, the relative labeling of myocardial ^13^C_3_-pyruvate (m+3) and [4,5-^13^C_2_]α-ketoglutarate (m+2) indicates the combined contribution of exogenous glucose and other carbohydrates entering the mitochondria via pyruvate dehydrogenase (V_PDH_), as a percentage of total substrate input (including fatty acids and ketones) to the TCA cycle via citrate synthase (V_CS_) ^26,27^. Since the pyruvate and α-ketoglutarate pools are small and difficult to analyze, we measured the enrichment of their reciprocal pools, namely ^13^C_3_-alanine (m+3) which rapidly equilibrates with ^13^C_3_-pyruvate (m+3), and [4,5-^13^C_2_]glutamate (m+2) which equilibrates with [4,5-^13^C_2_]α-ketoglutarate (m+2) (Figure 2, B and C) ^27^. The analysis of myocardial tissue from control mice by mass spectroscopy indicated that 37% of the total mitochondria oxidation (V_CS_) came from carbohydrate utilization (V_PDH_) in awake mice after a 6-hour fast (Figure 2, D and F). The V_PDH_/V_CS_ ratios were comparable in the left and right atria in control mice (Figure 2, D and F). However, atrial AMPK dKO mice showed a significantly increased V_PDH_/V_CS_ ratio in both atria as compared to littermate controls (Figure 2, D and F), indicating a shift in atrial mitochondrial metabolism towards pyruvate oxidation. The V_PDH_/V_CS_ ratios in the left and right ventricles were similar to those measured in atria in control mice but did not change in the atrial AMPK dKO mice, indicating that selective atrial AMPK deletion caused chamber-specific metabolic alterations (Figure 2, D and G).

Ketones are produced by the liver and their circulating plasma concentrations increase during periods of low food intake or prolonged exercise ^28^. The heart and other tissues readily take up ketones from the plasma and convert them to acetyl-CoA, which then enters the TCA cycle and is oxidized in the mitochondria for energy production. Although generally accounting for a low percentage (<10%) of ATP production ^25^, ventricular ketone metabolism is increased in heart failure and potentially also in AF ^7,29^. Thus, we studied the incorporation of ketones into the TCA cycle using LC-MS/MS methodology after infusion of [^13^C_4_]β-hydroxybutyrate in a separate cohort of mice. The ratio of [4,5-^13^C_2_]glutamate (m+2) to [^13^C_4_]β-hydroxybutyrate (m+4) was used to assess the relative contribution of β-hydroxybutyrate (V_BDH_) to total mitochondrial oxidation (V_CS_) (Figure 2, B and C). Control mice showed similar relative rates of V_BDH_ to total mitochondrial oxidation (V_BDH_/V_CS_) in the atria and ventricles, accounting for ∼5% of total mitochondria oxidation (Figure 2, E and F). Furthermore, no differences were detected between relative rates of β-hydroxybutyrate oxidation in either the atria or ventricles from atrial AMPK dKO mice compared to littermate controls (Figure 2, E and G).

The oxidation of fatty acids relative to total mitochondrial oxidation (V_FA_/V_CS_) was indirectly assessed by subtracting the myocardial V_BDH_/V_CS_ and V_PDH_/V_CS_ ratios from “1”, i.e. [1 - (V_BDH_/V_CS_ + V_PDH_/V_CS_)] (Figure 2C). Since this calculation does not take into account the minor contribution of amino acid oxidation, it slightly overestimates the contribution of fatty acid oxidation to overall mitochondrial oxidation. Using this approach, we calculated that fatty acid oxidation accounts for ∼60% of total atrial mitochondria oxidation in control mice (Figure 2F), consistent with previous measurements of myocardial substrate utilization ^25^. The loss of AMPK activity led to a significant reduction in the relative contribution of fatty acids to atrial TCA cycle flux, with V_FA_/V_CS_ accounting for only ∼30% of total mitochondrial oxidation in atrial AMPK dKO mice (Figure 2F). As anticipated, there was no difference in the contribution of fatty acids to total mitochondrial oxidation in ventricles in atrial AMPK dKO mice (Figure 2G). Thus, these results indicate that AMPK has an important homeostatic role in maintaining atrial fatty acid oxidation in awake mice studied under normal physiological conditions.

### Atrial AMPK deletion does not alter circulating metabolites or whole-body composition

To determine whether the shift in atrial metabolism might reflect in part a change in whole-body metabolism, we measured circulating plasma concentrations of glucose, non-esterified fatty acids, triglycerides, β-hydroxybutyrate, and insulin from control and atrial AMPK dKO mice. Although sarcolipin has some expression in skeletal muscle, there were no differences in these metabolic plasma parameters detected between the AMPK dKO and control groups, with the exception of a mild increase in β-hydroxybutyrate levels (Supplemental Figure 1B). Similarly, whole-body composition, food and water intake, energy expenditure, and respiratory exchange ratio (measured in metabolic cages) were also similar in the two groups (Supplementary Figure 1C). Thus, the changes that were observed in atrial metabolism in awake atrial AMPK dKO mice appeared to be attributed primarily to changes in the atrial myocardium, rather than substrate delivery.

### AMPK deletion increases atrial glucose uptake

Glucose metabolism plays a crucial role in maintaining myocardial energy homeostasis, especially during ischemia and heart failure ^30^. In order to better understand the mechanisms responsible for the increased atrial pyruvate oxidation observed in the atrial AMPK dKO mice, which is a reflection of both glucose and lactate oxidation, we assessed glucose uptake and phosphorylation *in vivo* after the administration of [^14^C]2-deoxyglucose in conscious mice, by measuring the accumulation of [^14^C]2-deoxyglucose-6-phosphate in the atrial tissue (Figure 2A). These experiments revealed that atrial glucose uptake was increased by 155% in the left atria and 82% in the right atria in the AMPK dKO mice as compared to control mice (Figure 3A). As anticipated, glucose uptake in the ventricles was similar in control and AMPK dKO mice (Figure 3A). The increased atrial glucose uptake and phosphorylation suggests that increased glycolysis contributes to the augmentation in pyruvate or carbohydrate oxidation (V_PDH_) that was observed in AMPK dKO mice.

**Figure 3.**
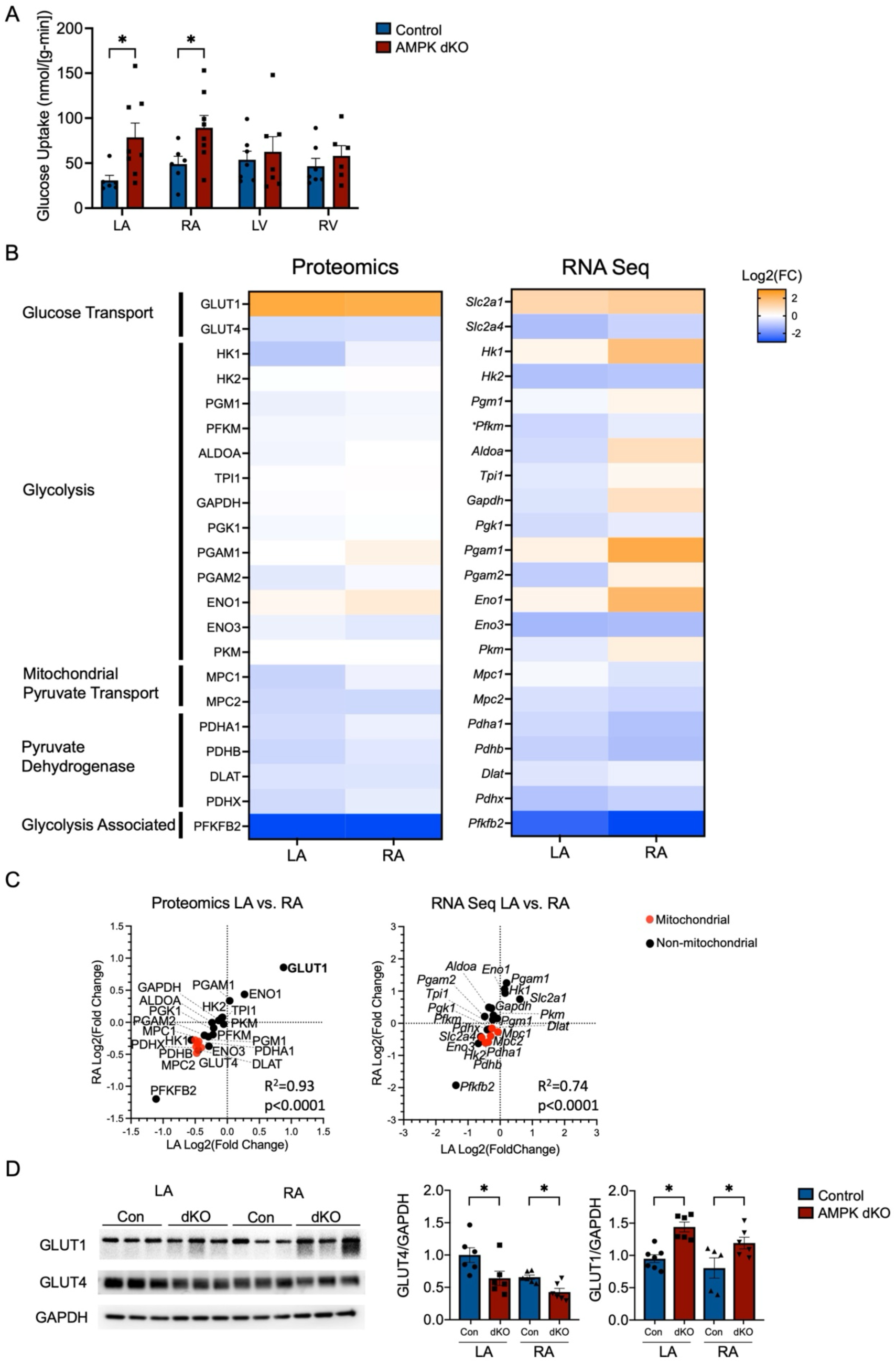
Atrial AMPK deletion increases atrial glucose uptake and glucose transporter GLUT1 protein expression. (A) Conscious mice were injected with [^14^C]2-deoxyglucose (intravenously) to assess glucose uptake in the left atria (LA), right atria (RA), left ventricle (LV), and right ventricle (RV) in control and atrial AMPK dKO mice at 4-6 weeks of age (n=6-8 per group). (B) Heatmap of proteomics and RNA-sequencing (RNA-seq) analyses of atrial tissue from control and AMPK dKO mice at 6 weeks of age (n=3 per group for both RNA-seq and proteomics), indicating changes in proteins and their corresponding RNA transcripts involved in glucose transport, glycolysis, and pyruvate entry into the mitochondria. Data show fold-changes (AMPK dKO vs. control) for LA and RA. (C) Comparison of proteomics and RNA-seq determined changes in glucose metabolism pathway components in the right and left atria of AMPK dKO vs. control mice. Proteins localized to mitochondria are indicated in red, and other proteins in black. (D) Representative immunoblots showing the atrial expression of glucose transporters GLUT1 and GLUT4 with quantification relative to GAPDH (n=3 per group). Data are shown as mean ± SEM; Significance determined by unpaired 2-tailed Student’s *t* test. *p < 0.05 AMPK dKO vs. control.

### Atrial AMPK alters expression of glucose transporters and glucose oxidation-related proteins

In order to elucidate whether components of the pathway responsible for glucose and fatty acid oxidation might have undergone metabolic reprogramming in the atrial AMPK dKO mice, we performed proteomic and bulk RNA-sequencing (RNA-seq) analyses on atrial myocardial tissue. The proteomics analysis identified 6609 unique proteins and demonstrated broad alterations in the expression of proteins involved in cellular metabolism (Supplemental Figure 2A). The RNA-Seq analysis yielded 52535 unique transcripts and, similarly, demonstrated alterations in transcripts for proteins involved in cellular metabolism (Supplemental Figure 2C).

To better understand the mechanisms contributing to increased glucose uptake, we first focused on the expression of the glucose transporters GLUT1 and GLUT4, which are the major glucose transporters in the heart ^31^. The former is responsible for basal glucose transport under fasting conditions associated with low levels of plasma insulin, while the latter mediates insulin-stimulated and ischemia-triggered glucose uptake in insulin-sensitive tissues including the heart^31^. The proteomic results indicated a significant increase in atrial GLUT1 expression, while there was a decrease in GLUT4 content in the atrial AMPK dKO mice (Figure 3, B and C). Subsequent immunoblotting of atrial homogenates confirmed the increase in GLUT1 and decrease in GLUT4 expression (Figure 3D). These studies were performed under fasting conditions, where GLUT1 is primarily responsible for heart glucose uptake with little contribution from GLUT4, and are concordant with the measurements showing increased deoxyglucose uptake.

The proteomic analysis also revealed variable changes in atrial glycolytic protein expression in the atrial AMPK dKO compared to control mice (Figure 3, B and C). Hexokinase 1 (HK1) expression was decreased in both atria, but hexokinase 2 (HK2), the more dominant isoform in the heart ^10,12^, displayed no significant change in expression (Figure 3, B and C). The muscle isoform of phosphofructokinase 1 (PFKM, also known as PFK1) also showed a decrease in expression in both atria while pyruvate kinase M (PKM) showed no change in expression (Figure 3, B and C). Interestingly, the expression of 6-phosphofructo-2-kinase/fructose-2,6-bisphosphatase 2 (PFKFB2, also known as PFK2), a key regulator of glycolysis, was downregulated in both atria (Figure 3, B and C). PFK2 is a bifunctional enzyme that both synthesizes and degrades fructose-2,6-bisphosphate (F2,6BP), an allosteric activator of PFK1, but the effect of this downregulation on glycolytic flux is not well established. Interestingly, the expression of mitochondrial proteins contributing to glucose oxidation was decreased in AMPK dKO atria, including mitochondrial pyruvate carriers (MPC1 and MPC2) and pyruvate dehydrogenase subunits (PDHA1, PDHB, DLAT, PDHX) (Figure 3, B and C). However, glycolytic enzymes and PDH are all strongly regulated by allosteric mechanisms, so that protein expression does not predict atrial glycolytic flux or the capacity for glucose oxidation.

The results of atrial RNA-seq analysis largely paralleled the proteomics data with regards to the glycolytic pathway, indicating that the observed proteomic changes were determined in part by transcriptional regulation (Figure 3, B and C). Notably, *Slc2a1* transcripts encoding GLUT1 were significantly increased in both atria, while *Slc2a4* transcripts encoding GLUT4 were decreased in the left atria, suggesting that transcriptional regulation plays a significant role in the observed changes in glucose transporter expression (Figure 3, B and C). Exceptions included an upregulation of HK1 in the right atrium and a downregulation of HK2 in both atria, contrasting with the proteomic findings.

### AMPK deletion reduces atrial long-chain fatty acyl-CoA content

Myocardial fatty acid oxidation is critical for myocardial energy production under normal physiological conditions. Fatty acids derived from both albumin-bound fatty acids and triglyceride in plasma are taken up across the sarcolemma and converted to fatty acyl-CoA intermediates prior to their entry into the mitochondria for oxidation. To assess the availability of fatty acyl-CoAs for mitochondrial metabolism, we used LC-MS/MS to measure the content of several myocardial C10 – C22 fatty acyl-CoA species. Both left and right atria from the AMPK dKO mice demonstrated an approximate 60% reduction in the content of long-chain fatty acyl-CoAs, including the acyl-CoA intermediates of palmitate, oleate, and linoleic acid, as compared to controls (Figure 4A); these latter fatty acids are major substrates for myocardial metabolism^32^. These findings highlight the important physiological role of AMPK in maintaining fatty acid availability for atrial mitochondrial oxidative metabolism.

**Figure 4:**
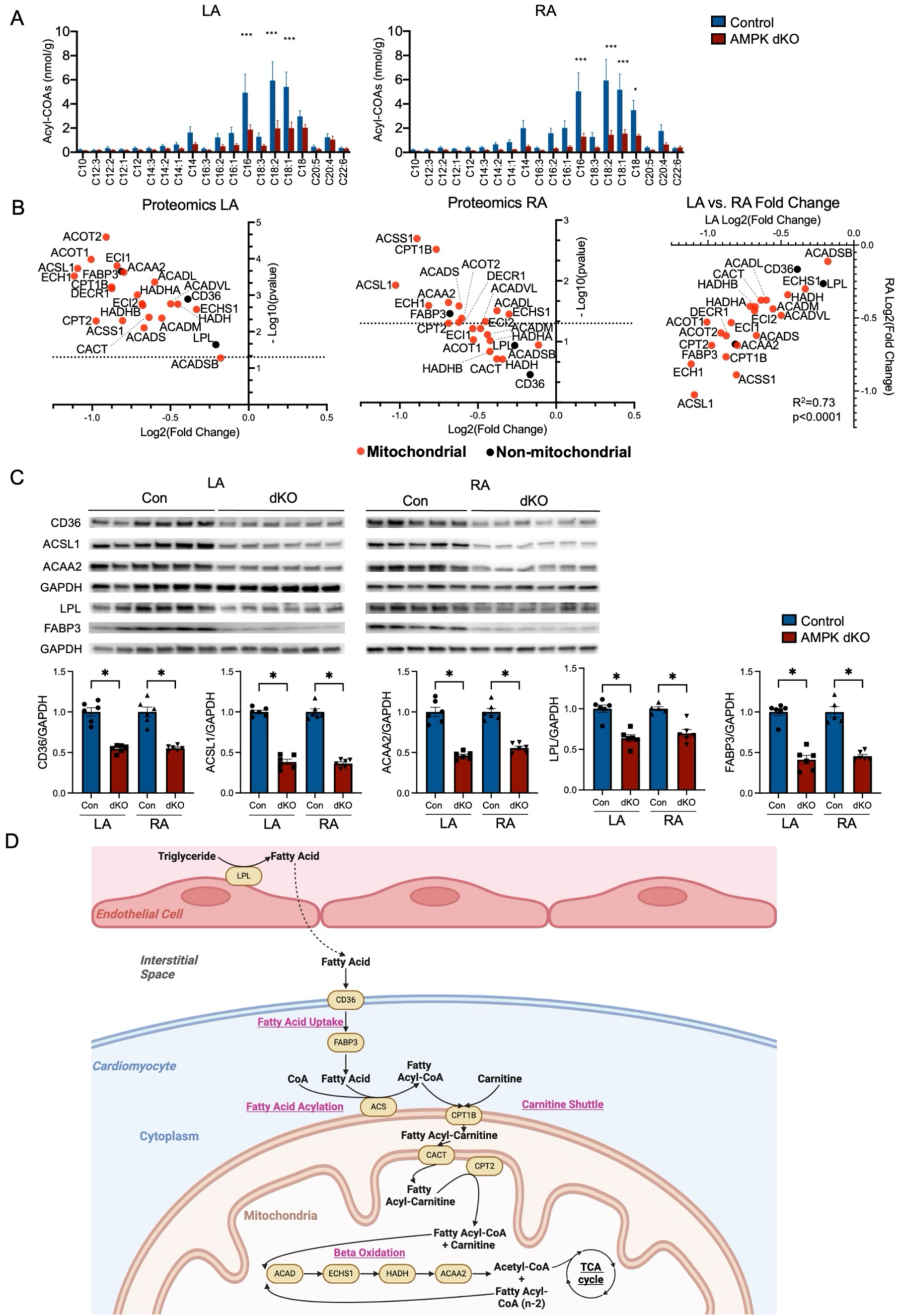
Atrial AMPK deletion reduces atrial long-chain fatty acyl-CoA content and downregulates the fatty uptake and oxidation pathways. (A) Content of myocardial long-chain fatty acyl-CoA species in the left atria (LA) and right atria (RA) in control and atrial AMPK dKO mice at 4 weeks of age measured by LC-MS/MS. Data are shown as mean ± SEM (n=8-9 per group). Significance determined by 2-way ANOVA with Sidak’s multiple-comparisons test. *p < 0.05, ***p<0.001 AMPK dKO vs. control. (B) Proteomic analysis of proteins responsible for fatty acid metabolism, comparing atrial AMPK dKO and control mice at 6 weeks (n=3 per group), for LA and RA. Data show fold-change (AMPK dKO vs. control), statistical significance between groups, and the correlation between left and right atrial changes. Proteins localized to mitochondria are indicated in red, and other proteins in black. Significance determined by unpaired 2-tailed Student’s t test. AMPK dKO vs. control. Graph includes dashed line that indicates p=0.05. (C) Representative immunoblots showing the expression of proteins responsible for fatty acid uptake (CD36, LPL and FABP3), and oxidation (ACSL1, ACAA2) with quantification relative to GAPDH in control and atrial AMPK dKO mice at 6 weeks of age (n=5-6 per group). Data are shown as mean ± SEM; Significance determined by unpaired 2-tailed Student’s *t* test. *p < 0.05 AMPK dKO vs. control (D) Schematic representation of proteomic results showing the downregulation of proteins (highlighted in beige ovals) involved in fatty acid uptake, acylation, mitochondrial entry, and beta oxidation in mice with atrial AMPK α_1_α_2_ deletion.

### AMPK deletion downregulates atrial fatty acid oxidation-related protein expression

Proteomics analysis of atrial tissue revealed a striking and consistent pattern of changes in the proteins associated with fatty acid metabolism, comparing the atrial AMPK dKO and control mice. The reduction in atrial long-chain fatty acyl-CoA levels, in the absence of a fall in circulating plasma fatty acid concentrations, raised the possibility of decreased cellular fatty acid uptake in the atrial AMPK dKO mice. The import of fatty acids from triglycerides and circulating lipoproteins, requires the action of lipoprotein lipase (LPL) on the endothelial surface. LPL is known to be essential for the hydrolysis of triglycerides, making long chain fatty acids available for cellular uptake. Triglycerides are an important source of fatty acids for the myocardium ^33^. Long-chain fatty acids then rely on the sarcolemma membrane transporter CD36 and the fatty acid binding protein FABP3 for entry into cardiomyocytes. The proteomic analysis indicated downregulation of LPL, CD36, and FABP3 in the atrial AMPK dKO mice: FABP3 levels decreased significantly in both atria, while LPL and CD36 exhibited significantly reduced content in the left atrium (Figure 4B). To confirm these findings, we selected a subset of key proteins involved in fatty acid uptake and oxidation for further validation. We performed immunoblotting, which demonstrated significant reductions in LPL, CD36, and FABP3 expression in both the left and right atria of the AMPK dKO mice (Figure 4C), confirming a substantial downregulation in proteins responsible for atrial fatty acid uptake.

In addition, the proteomic analysis revealed decreased expressions of key proteins involved in fatty acid oxidation in atrial AMPK dKO mice. These included ACSS1 and ACSL1 responsible for the acylation of short and long-chain fatty acids, respectively, as well as CPT1B, CACT, and CPT2, which mediate fatty-acyl transport into the mitochondria (Figure 4B). Immunoblots confirmed that there was significant downregulation of ACSL1 in both the left and right atria of the AMPK dKO mice (Figure 4C). There was also downregulation of the enzymes involved in mitochondrial beta-oxidation, including acyl-CoA dehydrogenases (ACADS, ACADM, ACADL, ACADVL, ACADSB), enoyl-CoA hydratases (ECHS1, ECH1, HADHA), enoyl-CoA isomerases (ECI1 and ECI2), 3-hydroxyacyl-CoA dehydrogenase (HADH, HADHA), and 3-ketoacyl-CoA thiolase (ACAA2, HADHB) (Figure 4B). Immunoblots also confirmed significant downregulation of ACAA2 in both atria of the AMPK dKO mice (Figure 3C). There was a good correlation (R^2^=0.71) between the proteomic alterations in the left and right atria, although the changes in the left atrium appeared to be more consistent and were more uniformly statistically significant. These latter results overall underscore the important role of AMPK in maintaining the expression of proteins crucial for mitochondrial fatty acid uptake and oxidation in the atria. A schematic representation of the proteomic results highlights the comprehensive downregulation of proteins involved in fatty acid uptake, acylation, carnitine shuttle, and beta-oxidation, providing a summary of the alterations observed (Figure 4D).

### AMPK regulates the PGC1-α/PPARα/RXR pathway in atrial cardiomyocytes

To further elucidate the mechanisms underlying the changes in fatty acid oxidation, we then focused on transcriptional regulation of fatty acid oxidation-related protein genes. In striated muscle, this process is regulated by the peroxisome proliferator-activated receptor alpha (PPARα) and retinoid X receptor (RXR) in complex with peroxisome proliferator-activated receptor gamma coactivator 1-alpha (PGC1-α) ^34-39^. Although AMPK is known to regulate PGC-1α in skeletal muscle, evidence establishing a clear link in heart is lacking. In order to assess the extent to which atrial AMPK regulates PGC1-α, PPARα, and RXR, we first analyzed the RNA-seq data for PGC1-α, PPARα, and RXR. The analysis demonstrated downregulation of both left and right atrial transcripts encoding PGC1-α (*Ppargca1*) and PPARα (*Ppara*) in AMPK dKO mice (Supplemental Figure 3). Interestingly, while the RNA-seq analysis demonstrated no significant change for RXRα transcripts (*Rxra*), we did observe significant downregulation of *Rxrg* transcripts encoding RXRγ, which might also contribute to the regulation of the PPARα/PGC1α pathway (Supplemental Figure 3). Immunoblots confirmed a significant decrease in the expression of PGC1-α in both atria of AMPK dKO mice (Figure 5A). We were unable to identify an antibody that specifically detected PPARα or RXR expression on immunoblots; these transcription factors are expressed in relatively low abundance and were also not detected in the proteomic analysis. ERR has also been shown to increase the transcription of genes encoding proteins responsible for fatty acid oxidation in the left ventricle ^36^. The RNA-seq analysis showed no evident change in *Esrra* or *Esrrg* transcripts, encoding ERRα and ERRγ, respectively, but *Esrrb* transcripts encoding the low abundance ERRβ were increased in the right atrium (Supplemental Figure 3).

**Figure 5.**
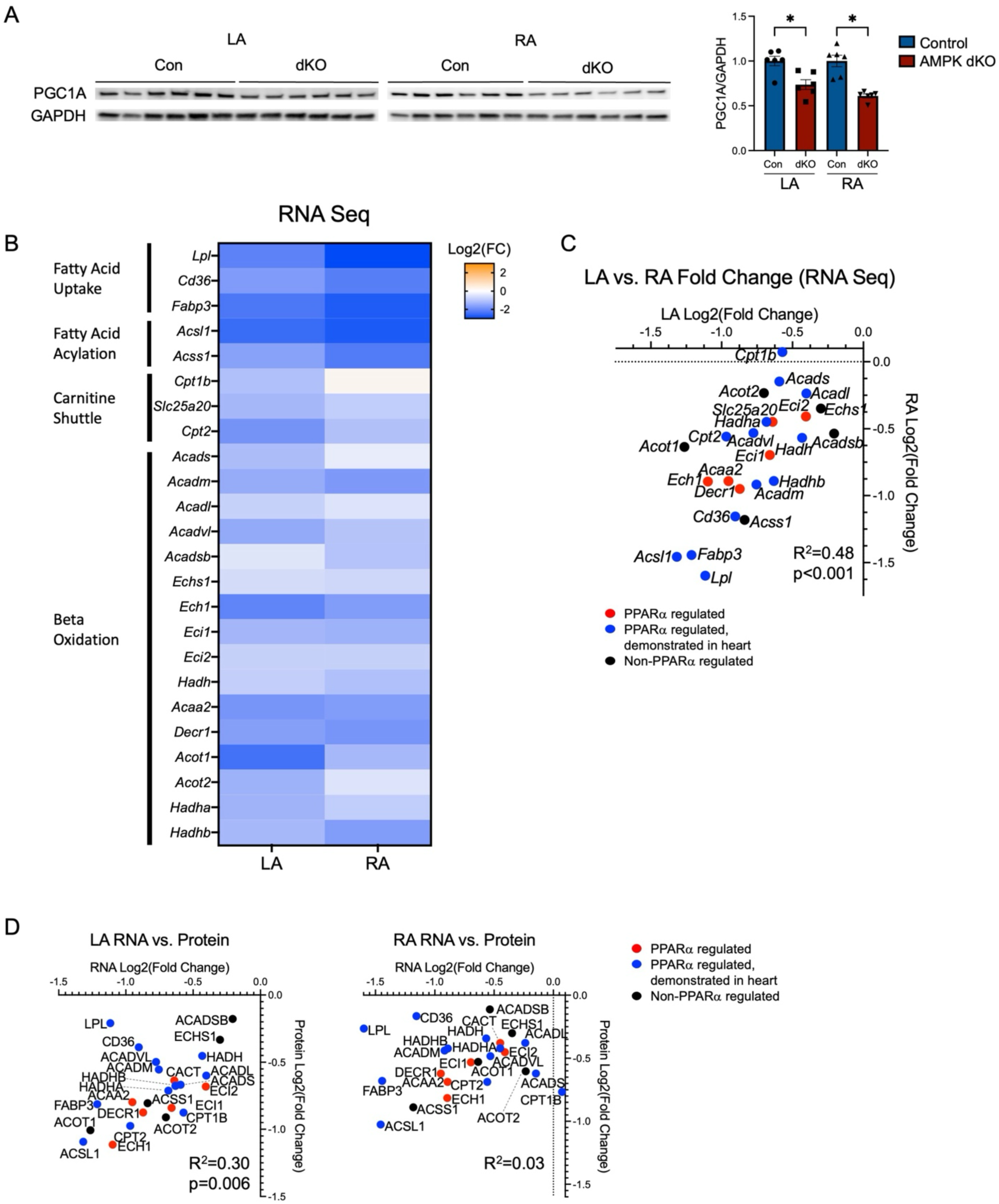
Atrial AMPK deletion downregulates PGC-1α/PPARα/RXR regulated genes critical to atrial fatty acid uptake and oxidation. (A) Representative immunoblots showing the expression of PGC-1α with quantification relative to GAPDH in left atria (LA), right atria (RA) of control and atrial AMPK dKO mice at 6 weeks of age (n=6 per group). (B) Heatmap of RNA-sequencing (RNA-seq) analysis of transcripts encoding fatty acid metabolism proteins comparing atrial control and AMPK dKO mice at 6 weeks. Data show fold-change (AMPK dKO/Control) for LA and RA. (C) Quantitative presentation of alterations in transcripts encoding proteins for fatty acid metabolism for LA and RA showing the correlation between left and right atria changes (n=3 per group). Data includes genes known to be PPARα regulated in heart (in blue), in other tissues (in red) or non-PPARα regulated (in black). (D) Correlation between the degree of downregulation of RNA transcripts (RNA-seq) and protein expression (proteomic analysis) of components of the fatty acid metabolic pathway, comparing control and AMPK dKO mice, for LA and RA. Data includes genes known to be PPARα regulated in heart (in blue), in other tissues (in red) or non-PPARα regulated (in black). Data are shown as mean ± SEM; Significance determined by unpaired 2-tailed Student’s *t* test. *p < 0.05 AMPK dKO vs. control.

In order to assess the extent to which atrial AMPK regulates PPAR-α mediated gene transcription, we then examined the RNA-seq data for fatty acid oxidation-related protein transcripts. We observed that the results of RNA-seq analysis mirrored the proteomics data. Specifically, AMPK dKO mice demonstrated decreases in atrial transcripts encoding proteins involved in fatty acid uptake (*Lpl*, *Cd36*, *Fabp3*), fatty acid acylation (*Acss1*, *Acsl*1), carnitine shuttle (*Cpt1b*, *Slc25a20*, *Cpt2*), and fatty acid oxidation (*Acads*, *Acadm*, *Acadl*, *Acadvl*, *Acadsb*, *Echs1*, *Ech1*, *Eci1*, *Eci2*, *Decr1*, *Hadh*, *Hadha*, *Hadhb*, *Acaa2*, *Acot1*, *Acot2*) (Figure 5, B and C). Several of these genes are known to be PPAR-α regulated in the heart or other tissues, including genes responsible for fatty acid uptake (*Lpl*, *Cd36*, *Fabp3*), fatty acid acylation (*Acsl1*), and fatty acid beta-oxidation enzymes (*Acaa2*). Furthermore, there was a reasonably good correlation between the extent of downregulation in the left compared to the right atria of AMPK dKO hearts (R^2^=0.48) (Figure 5C). To determine whether the decrease in the content of mRNA transcripts paralleled the degree of downregulation in the expression of corresponding proteins, we compared the results of RNA-seq and proteomic analysis in AMPK dKO and control hearts. There was a positive correlation between the downregulation of atrial fatty acid metabolism proteins and their respective transcripts, indicating the importance of transcriptional regulation in metabolic reprogramming in the atrial AMPK dKO mice (Figure 5D). Notably, metabolic proteins that are known to be encoded by PPARα regulated genes, exhibited a greater degree of downregulation of protein expression and transcript content in the AMPK dKO vs. control mice in both atria.

Thus, taken together these results indicate that loss of AMPK impairs PGC1-α/PPARα/RXR expression in the atria and establishes a link between AMPK and the PGC1-α/PPARα/RXR pathway, which is critical to maintaining atrial fatty acid cellular uptake and entry into the mitochondria for TCA cycle oxidation.

## Discussion

The findings of this study demonstrate that AMPK has a critical homeostatic role in regulating atrial metabolism. Specifically, mice with atrial deletion of the AMPK α1 and α2 catalytic subunits demonstrated impaired atrial mitochondrial fatty acid oxidation, associated with increased glucose uptake and a shift towards greater carbohydrate oxidation *in vivo.* The impaired utilization of fatty acids was paralleled by a reduction in the atrial content of myocardial long-chain fatty-acyl CoA intermediates. This reduced availability of intracellular fatty acids for mitochondrial oxidation was associated with a downregulation of proteins responsible for cellular fatty acid uptake and acylation. In addition, multiple proteins mediating subsequent mitochondria fatty acid entry and oxidation were also downregulated in the AMPK dKO atria. This metabolic remodeling was accompanied by decreased expression of PGC1-α, and reduced content of mRNA transcripts encoding PPARα, PGC1-α, and RXR with transcriptional downregulation of multiple PPARα-regulated genes encoding proteins responsible for fatty acid metabolism. In a presumably compensatory manner, there was augmented carbohydrate oxidation as indicated by an increased V_PDH_/V_CS,_ which reflects the combined contributions of glucose, lactate and pyruvate to mitochondrial oxidative metabolism. This increase in V_PDH_/V_CS_ was accompanied by enhanced atrial deoxyglucose uptake indicating augmented glucose transport and phosphorylation *in vivo*, and increased expression of the glucose transporter GLUT1 in atrial tissue from the AMPK dKO mice. These results indicate that major metabolic reprogramming develops consequent to the loss of atrial AMPK activity and implicates a link between AMPK and PGC1-α, PPARα and RXR in the atria, overall highlighting the important homeostatic role of AMPK in regulating atrial substrate metabolism.

This investigation characterizes, for the first time to our knowledge, atrial substrate utilization and oxidation *in vivo*. Using a combination of LC-MS/MS methodology combined with infusions of ^13^C -labeled substrates, we measured the relative rates of atrial pyruvate, ketone and fatty acid oxidation. The studies were performed in conscious mice to avoid the confounding and important hemodynamic and metabolic effects of anesthesia on *in vivo* metabolism ^40-42^. As such, they provide novel physiological insight into the regulation of atrial metabolism in the context of an intact *in vivo* metabolic, hormonal and hemodynamic milieu, that is difficult to replicate *in vitro*. The relative rates of pyruvate oxidation to total mitochondrial oxidation (V_PDH_/V_CS_) and ketone oxidation to total mitochondrial oxidation (V_BDH_/V_CS_) were measured from the steady-state labeling of [4,5-^13^C_2_]glutamate (m+2) during continuous intravenous infusions of [^13^C_6_]glucose and [^13^C_4_]β-hydroxybutyrate, respectively ^27^. Fatty acid oxidation was assessed indirectly from the remaining unlabeled fraction of glutamate, [1 - (V_BDH_/V_CS_ + V_PDH_/V_CS_)]. It should be noted that this approach provides an integrated assessment of the contributions of both exogenous and endogenous substrate to oxidative metabolism in the TCA cycle. While the calculation of fatty acid oxidation is over-estimated due to a small component of amino acid oxidation, primarily from branch chain amino acids, this contribution is quantitatively minor compared to fatty acid oxidation.

Using this LC-MS/MS technique, we found that atrial AMPK dKO mice demonstrated more than a 50% reduction in the relative contribution of fatty acids to mitochondrial oxidative metabolism. Interestingly, there was also a similar 50% decrease in the atrial content of long-chain fatty acyl-CoA, which are the major substrate for mitochondrial fatty acid oxidation. These results indicated a potential defect in cellular fatty acid uptake and/or acetylation. Consistent with this observation, multiple proteins mediating cellular fatty acid entry (LPL, CD36, FABP3) and fatty acid acylation (ACSL1, ACSS1), were downregulated in the AMPK dKO atria. Furthermore, in addition to the lack of fatty acyl-CoA substrate available for mitochondrial oxidation, we also identified downregulation of proteins responsible for mitochondrial fatty acid import and most of the proteins involved in fatty acid oxidation in the AMPK dKO atria. These results together indicate that AMPK is critical to maintaining both atrial fatty acid substrate supply and mitochondrial fatty acid oxidation, such that the lack of AMPK results in impaired atrial physiological fatty acid oxidation in conscious AMPK dKO mice.

The expression of gene transcripts encoding the proteins in the fatty acid metabolism pathway were also found to be downregulated in the AMPK dKO atria, based on the RNA-seq analysis. The extent of downregulation of mRNA transcripts was highly correlated with changes in proteins, based on the proteomic analysis, in both the left and right atria. Several of these genes are known to be PPARα regulated in either the heart or other tissues, including those encoding LPL, CD36, FABP3, ACSL1, and ACAA2, as well as multiple proteins involved in beta-oxidation ^38,39^. These findings indicate that PPARα has a parallel and critical role in regulating atrial fatty acid oxidation and lipid homeostasis as it does in the ventricle ^43,44^. We were unable to identify an antibody that specifically detected PPARα to further quantify PPARα expression on immunoblots. However, PPARα activity is also modulated by RXR and the concentration of intracellular fatty acids (endogenous ligands for PPARα), so that the downregulation of RXR and the low content of fatty-acyl CoAs would also be expected to contribute to the diminished PPARα-mediated gene regulation observed in the AMPK dKO atria.

Our results also provide evidence that AMPK regulates PGC-1α expression in the atria. Although the link between AMPK and PGC-1α is well-established in skeletal muscle, prior data supporting this link in the heart are limited. Loss of AMPK activity was shown to have little effect on baseline PGC-1α in the normal left ventricle, although lack of AMPK did blunt PGC-1α activation in the context of myocardial ischemia ^45^. No prior studies to our knowledge have explored this mechanism in the atria. We found that PGC-1α mRNA transcripts and protein expression were diminished in both left and right atria from AMPK dKO hearts. Furthermore, the activity of PGC-1α is known to be post-transcriptionally regulated. In skeletal muscle, AMPK increases NAD^+^ concentrations, activating sirtuin-1, which deacetylates and activates PGC-1α ^46^. AMPK has also been reported to directly phosphorylate and activate PGC-1α at Thr^177^ and Ser^538^, which then increases its interaction with PPAR-α and RXR to augment the transcription of genes involved in mitochondrial fatty acid oxidation ^34-39^. Loss of these additional mechanisms of PGC-1α activation in atria would be predicted to further impair PGC-1α action in the AMPK dKO atria.

In the context of diminished fatty acid oxidation, the atrial AMPK dKO mice appeared to compensate with an upregulation of atrial carbohydrate oxidation as reflected in the V_PDH_/V_CS_ ratio. The V_PDH_/V_CS_ index provides an integrated assessment of the contribution of carbohydrates to oxidative metabolism *in vivo*. These carbohydrates include exogenous glucose, lactate and pyruvate, as well as endogenous glucose derived from glycogen, endogenous lactate and pyruvate derived from glucose and to a lesser extent from alanine. In addition, enhanced exogenous glucose uptake was also observed in the atria of AMPK dKO mice, as indicated by increased [^14^C]2-deoxyglucose accumulation. This increased glucose uptake was associated with, and likely attributed to, an increase in the atrial expression of the GLUT1 transporter. Similar up-regulation of GLUT1 has also been observed as an adaptive response of the left ventricle to metabolic stress in heart failure, which mimics a more fetal metabolic pattern characterized by increased glucose oxidation and reduced fatty acid oxidation ^23,24^. Interestingly, the atrial expression of the insulin-regulated GLUT4 transporter was decreased in the atrial AMPK dKO mice. GLUT4 activity is highly regulated by its subcellular location in striated muscle, and GLUT4 translocation to the sarcolemma is increased by insulin and ischemia ^31^. Under fasting conditions, there is typically little GLUT4 in the sarcolemma and thus the downregulation of GLUT4 is unlikely to have had impact on glucose transport in these experiments. In contrast, the GLUT4 downregulation would be expected to compromise glucose uptake in the fed state, when insulin levels are higher, which would predict that post-prandial atrial glucose uptake might also be compromised in these mice.

Ketones have emerged as an alternate substrate for the heart, particularly in the left ventricle in the context of heart failure and during treatment with SGLT2 inhibitors ^47^. We found that the contribution of ketones to mitochondrial oxidation (V_BDH_/V_CS_) in the atria was comparable (∼5%) to the ventricles of the conscious fasting control mice. However, the awake AMPK dKO mice displayed no evident change in atrial ketone oxidation, despite a slight upregulation in circulating β-hydroxybutyrate.

The substantial atrial metabolic remodeling that develops in these mice consequent to AMPK deletion potentially points to a critical physiological role of AMPK in the atria, in contrast to the ventricles, where AMPK inactivation has modest impact on the homeostasis of myocardial metabolism in the normal heart. AMPK deficient mouse models demonstrate relatively normal baseline myocardial left ventricular substrate utilization ^17,18,48^, with alterations in metabolism developing only under cardiac stress, such as ischemia ^17^. In this current study, we were unable to directly compare the effects of AMPK deletion in the atria and the ventricles, since we specifically utilized an atrial specific model to avoid potentially confounding effects of changes in ventricular function on the atria.

We recently demonstrated that this model of atrial AMPK deletion leads to early electrophysiological abnormalities with late structural remodeling and fibrosis, culminating in spontaneous atrial fibrillation ^22^. In order to elucidate the primary impact of AMPK deletion on atrial metabolism, while avoiding the secondary effects of atrial remodeling and fibrillation, we performed the current experiments in young AMPK dKO mice at 4-6 weeks of age. These mice had indicators of early atrial arrhythmia with atrial premature complexes and short runs of atrial tachycardia, but not persistent atrial fibrillation. Thus, the results demonstrate that substantial reprogramming of cardiac metabolism develops early in these mice, which suggests that the metabolic alterations represent one of the underpinnings, rather than a consequence of atrial fibrillation.

One limitation of this study is that it does not define the extent to which metabolic reprogramming has a pathophysiological role in the electrophysiological remodeling, fibrosis and ultimate development of atrial fibrillation in AMPK dKO mice. Additional studies would be required to determine whether correction of the metabolic abnormalities, with strategies that activate PGC1-α, PPARα, or provide alternate substrates (ketone supplementation, SGLT2 inhibitors, or short and/or medium chain fatty acids), might mitigate the development of atrial remodeling and atrial fibrillation in this model. Nonetheless, the results provide an integrated insight into the metabolic reprograming that ensues from loss of atrial AMPK and elucidate the important homeostatic role of AMPK in maintaining physiological atrial metabolism *in vivo*.

## Acknowledgements

This research was funded in part by grants from the NIH: R01HL148008 (L.H.Y., F.G.A.), R01HL128069 (L.H.Y.), HL149344 (F.G.A.), R01DL133143 (G.I.S.), P30DK045735 (G.I.S.), R00 HL150234 (L.G.). We would also like to thank Dr. Stephen Young for providing anti-LPL mouse IgG antibody.

## Methods

### Experimental animals

Atrial AMPK α1 and α2 double knockout mice (AMPK dKO) were generated by breading the homozygous AMPK α^flox/flox^α2^flox/flox^ mice with C57BL/6 mice expressing Cre-recombinase under the control of the sarcolipin (*Sln*) promoter, as previously described ^49^. Sarcolipin Cre AMPK α^flox/flox^α2^flox/flox^ mice and littermate controls without Cre-recombinase were used for all studies. All mice were on a pure C57BL/6 background and kept under constant temperature and humidity in a 12-hour controlled light/dark cycle (7:00am-7:00pm). Mice were multiply housed (3-4/standard cage) and fed a standard chow diet. Only male mice were used in this study. Assessment of food and water intake, and indirect calorimetry were performed using Columbus Lab Animal Monitoring System metabolic cages (Columbus Instruments, Columbus OH). Jugular venous polyethylene catheters were surgically placed one week prior to the tracer infusions and housed in single cages. Only mice that recovered pre-surgery body weight were studied. All animal experiments were approved by the Yale Animal Care and Use Committee.

### Metabolic flux studies

After a 6-hour fast, a primed (3 mg/kg-min for 5 min) continuous infusion of [^13^C_6_]glucose (1 mg/kg-min for 120 minutes) was administered to measure the relative rates of cardiac pyruvate oxidation (V_PDH_) to total mitochondrial oxidation (V_CS_), adapting a method developed in skeletal muscle ^50^. To assess glucose uptake in the heart, mice were intravenously injected with [^14^C]2-deoxyglucose (10 □Ci) during the last 20 minutes of the [^13^C_6_]glucose infusions (see below). Whole blood was obtained by tail vein at 0, 90, 100, 110, and 120 minutes and immediately centrifuged for the assessment of plasma metabolites (see below). Mice were then anesthetized by an intravenous infusion of pentobarbital and the heart immediately excised, left and right atria freeze-clamped in aluminum tongs pre-chilled in liquid N_2_ and stored at -80°C for subsequent analyses. Frozen atrial and ventricular tissue was pulverized on dry ice and homogenates prepared for analysis by LC-MS/MS and GC/MS to determine the enrichments of [4,5-^13^C_2_]glutamate (m+2) and [^13^C_3_]alanine (m+3), respectively, for the calculation of myocardial V_PDH_/V_CS_ flux (Eq. 1) as previously described ^27,50^:

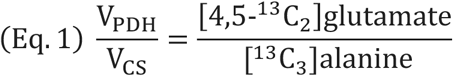

The ratio of [4,5-^13^C_2_]glutamate (m+2) and [^13^C_3_]alanine (m+3) reflects carbohydrate flux through V_PDH_ relative to V_CS_ (V_PDH_/V_CS_), such that a ratio of one indicates 100% pyruvate oxidation and any ratio less than one reflects dilution of the tracer from unlabeled acetyl-CoA (e.g. oxidation of fatty acids, ketones and/or ketogenic amino acids). Although true V_PDH_/V_CS_ flux requires measurement of [^13^C_2_]acetyl-CoA (m+2) and [^13^C_3_]pyruvate (m+3), these metabolites are difficult to measure reliably due to small pool sizes ^51,52^. We have previously shown that under steady-state conditions, intracellular [^13^C_3_]pyruvate (m+3) equilibrates with [^13^C_3_]alanine (m+3) and intracellular [^13^C_2_]acetyl-CoA (m+2) equilibrates with [4,5-^13^C_2_]glutamate (m+2) in the muscles of fasted rodents ^24^.

The relative rates of cardiac β-hydroxybutyrate oxidation (V_BDH_) to total mitochondrial oxidation (V_CS_) were measured in a paralleled cohort of mice following a primed (0.6 mg/kg-min x 5 min) continuous infusion (0.3 mg/kg-min x 120 min) of [^13^C_4_] β-hydroxybutyrate. Whole blood was obtained by tail vein at 0, 100, 110 and 120 minutes of the infusion, and processed for assessment of plasma metabolites. Left and right atria freeze-clamped, and homogenates were prepared as described above and analyzed by LC-MS/MS and GC/MS to determine enrichments. Myocardial V_BDH_/V_CS_ flux was measured using the following equation (Eq. 2):

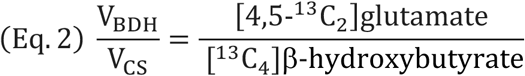

The ratio of [4,5-^13^C_2_]glutamate (m+2) to [^13^C_4_]β-hydroxybutyrate reflects ketone flux of β-hydroxybutyrate (V_BDH_) relative to V_CS_ (V_BDH_/V_CS_). Although true V_BDH_/V_CS_ flux requires measurement of [^13^C_2_]acetyl-CoA (m+2), [4,5-^13^C_2_]glutamate (m+2) was used as a surrogate for [^13^C_2_]acetyl-CoA (m+2) due to the aforementioned difficulties in reliably measuring isotopologues of acetyl-CoA. Myocardial β-hydroxybutyrate enrichment was measured by GC/MS ^53^. Myocardial glutamate enrichment was measured by LC-MS/MS ^24^.

The relative oxidation rate of fatty acids and ketogenic amino acids (V_FA +AA_) to total mitochondrial oxidation (V_CS_) was indirectly assessed from relative rates of myocardial V_BDH_/V_CS_ and V_PDH_/V_CS_ obtained during the [^13^C_6_]glucose and [^13^C_4_]β-hydroxybutyrate infusions (Eq. 3):

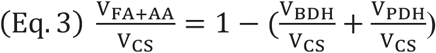

### Tissue glucose uptake measurements

Glucose uptake was measured in the heart following the bolus injection of [^14^C]2-deoxyglucose. Atrial and ventricular tissue were separately processed and [^14^C]2-deoxyglucose was separated from [^14^C]2-deoxyglucose using column chromatography to determine glucose uptake by comparing the plasma [^14^C] specific activity decay curve to tissue [^14^C] specific activity, as previously described ^27^.

### Plasma metabolite analyses

Blood samples were collected by tail vein after a 6-hour fast. Blood was immediately placed in heparinized-lithium tubes, separated by centrifugation, and plasma was stored at –80°C. Plasma glucose concentrations were measured using a YSI Glucose Analyzer (Yellow Springs, OH) and plasma insulin by ELISA (Mercodia). Plasma NEFAs were measured using kits from Wako Diagnostics, according to the manufacturer’s instructions. Plasma β-hydroxybutyrate was measured by COBAS.

### LC/MS/MS analysis of long-chain acyl-CoAs

Long-chain acyl-CoAs were extracted from frozen tissue samples (∼10-15 µg) and purified using a solid phase extraction method described previously ^54,55^ with minor modifications for desalting. A known amount of heptadecanoyl-CoA was added as an internal standard. OPC columns (Applied Biosystems, Foster City, CA) were used for solid phase extraction. Samples were dissolved in 10 µl of methanol/H2O for LC/MS/MS analysis.

A benchtop tandem mass spectrometer, API 6500 QTRAP (SCIEX), interfaced with a TurboIonspray ionization source or atmospheric pressure chemical ionization source was used. Peripherals included a Prominence UFLCXR HPLC system (Shimadzu). Long-chain acyl-CoAs were ionized in negative electrospray mode. Doubly charged ions and corresponding product ions were chosen as transition pairs for each CoA species (C16:1, C16:0, C18:2, C18:1, and C18:0) for selective reactions monitoring (SRM) quantitation. Total long-chain acyl-CoAs content was obtained from the sum of individual species. Methanol/H2O (60/40) was used as continuous flow at 300 µl/min, and 5 µl of sample was injected for analysis.

### RNA-sequencing

Total RNA quality was determined by estimating the A260/A280 and A260/A230 ratios by nanodrop. RNA integrity was determined by running an Agilent Bioanalyzer gel, which measures the ratio of the ribosomal peaks. Samples with RIN values of 5 or greater were recommended for library prep.

Using the Kapa RNA HyperPrep Kit with RiboErase (KR1351), rRNA was depleted starting from 25-1000 ng of total RNA by hybridization of rRNA to complementary DNA oligonucleotides, followed by treatment with RNase H and DNase to remove rRNA duplexed to DNA. Samples were then fragmented using heat and magnesium. 1st strand synthesis was performed using random priming. 2nd strand synthesis incorporated dUTPs into the 2nd strand cDNA. Adapters were then ligated, and the library was amplified. Strands marked with dUTPs were not amplified allowing for strand-specific sequencing. Indexed libraries that met appropriate cut-offs for both quantity and quality were quantified by qRT-PCR using a commercially available kit (KAPA Biosystems) and insert size distribution determined with the LabChip GX or Agilent Bioanalyzer. Samples with a yield of ≥0.5 ng/µl were used for sequencing.

Sample concentrations were normalized to 1.2 nM and loaded onto an Illumina NovaSeq flow cell at a concentration that yields 40 million passing filter clusters per sample. Samples were sequenced using 100bp paired-end sequencing on an Illumina NovaSeq according to Illumina protocols. The 10bp unique dual index was read during additional sequencing reads that automatically follow the completion of read 1. Data generated during sequencing runs were simultaneously transferred to the YCGA high-performance computing cluster. A positive control (prepared bacteriophage Phi X library) provided by Illumina was spiked into every lane at a concentration of 0.3% to monitor sequencing quality in real time.

Signal intensities were converted to individual base calls during a run using the system’s Real Time Analysis (RTA) software. Base calls were transferred from the machine’s dedicated personal computer to the Yale High Performance Computing cluster via a 1 Gigabit network mount for downstream analysis. Primary analysis - sample de-multiplexing and alignment to the human genome was performed using Illumina’s CASAVA 1.8.2 software suite. The data were used if the sample error rate was less than 2% and the distribution of reads per sample in a lane was within reasonable tolerance. Data were retained on the cluster for at least 6 months, after which the data were transferred to a tape backup system.

### Quantitative Proteomics

Atrial issues were pulverized in liquid nitrogen and lysed in buffer containing 10 M urea and the cOmplete™ protease inhibitor cocktail (Roche, #11697498001) by sonication at 4°C for 10 minutes (with 5 seconds on/off cycles) using a Cole-Parmer Ultrasonic Processor. The lysed samples were centrifuged at 20,000 x g for 1 hour to remove insoluble material. Atrial protein (700 µg) underwent reduction and alkylation using 10 mM dithiothreitol for 1 hour at 56°C, followed by 20 mM iodoacetamide in darkness for 45 minutes at room temperature. The samples were then diluted with 100 mM NH_4_HCO_3_ and digested with trypsin (Promega) at a ratio of 1:20 (w/w) overnight at 37°C. The digested peptides were purified using a C18 column (MacroSpin Columns, NEST Group INC). About 1 µg of the peptides were utilized for total proteome analysis.

The peptide samples were measured by data-independent acquisition (DIA) MS method as described previously ^24,56,57^, on an Orbitrap Fusion Tribrid mass spectrometer (Thermo Scientific) coupled to a nanoelectrospray ion source (NanoFlex, Thermo Scientific) and an EASY-nLC 1200 system (Thermo Scientific, San Jose, CA). A 150-min gradient was used for the data acquisition at a flow rate of 300 nL/min with the column temperature controlled at 60 °C using a column oven (PRSO-V1, Sonation GmbH, Biberach, Germany). The DIA-MS method consisted of one MS1 scan and 33 MS2 scans of variable isolated windows with 1 m/z overlapping between windows. The MS1 scan range was 350 – 1650 m/z and the MS1 resolution was 120,000 at m/z 200. The MS1 full scan AGC target value was set to be 2E6 and the maximum injection time was 100 ms. The MS2 resolution was set to 30,000 at m/z 200 with the MS2 scan range 200 – 1800 m/z and the normalized HCD collision energy was 28%. The MS2 AGC was set to be 1.5E6 and the maximum injection time was 50 ms. The default peptide charge state was set to 2. Both MS1 and MS2 spectra were recorded in profile mode. DIA-MS data analysis was performed using Spectronaut v18 ^58-60^ with directDIA algorithm by searching against the SwissProt mouse fasta file. The oxidation at methionine was set as variable modification, whereas carbamidomethylation at cysteine was set as fixed modification. Both peptide and protein FDR cutoffs (Qvalue) were controlled below 1% and the resulting quantitative data matrix were exported from Spectronaut. All the other settings in Spectronaut were kept as Default.

### Western blotting

Heart lysates were prepared in RIPA buffer with freshly added protease inhibitors (cOmplete MINI, Roche) and phosphatase inhibitors (PhosSTOP, Roche). After normalizing for equal protein concentrations by the BCA assay (Pierce), tissue homogenates were resuspended in Laemmli buffer sample buffer containing 4% 2-mercaptoethanol and separated on 4-8% Tris-Glycine Gels (Novex). Following a 1-hour semi-try transfer onto PVDF membranes, the membranes were blocked with 5% BSA (w/v) in TBST and probed with primary antibodies overnight at 4°C (listed below). After washing in TBST, membranes were incubated for 1-hour at room temperature with HRP-conjugated secondary antibodies (1:2000) diluted in blocking buffer. Bands were visualized by enhanced chemiluminescence (Pierce), and densitometry analysis was carried out using ImageJ software (NIH).

Cell Signaling Technology antibodies used were: phospho-ACC S79 (#3661, 1:1000 dilution), ACC (#3662, 1:1000 dilution), phospho-raptor S792 (#2083, 1:1000 dilution), raptor (#2280, 1:1000 dilution), AMPKα1/α2 (#2532, 1:1000 dilution), GAPDH (#2118, 1:1000 dilution), CD36 (#74002, 1:1000 dilution), ACSL1 (#9189, 1:1000 dilution). Abcam antibodies used were ACAA2 (ab314025, 1:1000 dilution) and PGC1α (ab191838, 1:1000 dilution). GLUT1 antibody was from Sigma-Aldrich (sab4200519, 1:1000 dilution) and GLUT4 antibody was from Santa Cruz Biotechnology (#7938, 1:500 dilution). LPL antibody was provided by Dr. Stephen Young.

### Statistics

All the figures were shown as mean ± SEM. For single-factorial analyses, an unpaired 2-tailed Student’s t test was performed to measure the differences in individual groups. One-way ANOVA with Tukey’s multiple-comparisons test were used to determine the statistical significance between 2 or more independent groups. Two-way ANOVA with Tukey’s multiple-comparisons test were used for results analysis. A P value of less than 0.05 was considered statistically significant. Data analyses were performed using GraphPad Prism software Version 9 (Graphpad, San Diego, CA).

## Figures

**Supplementary Figure 1.**
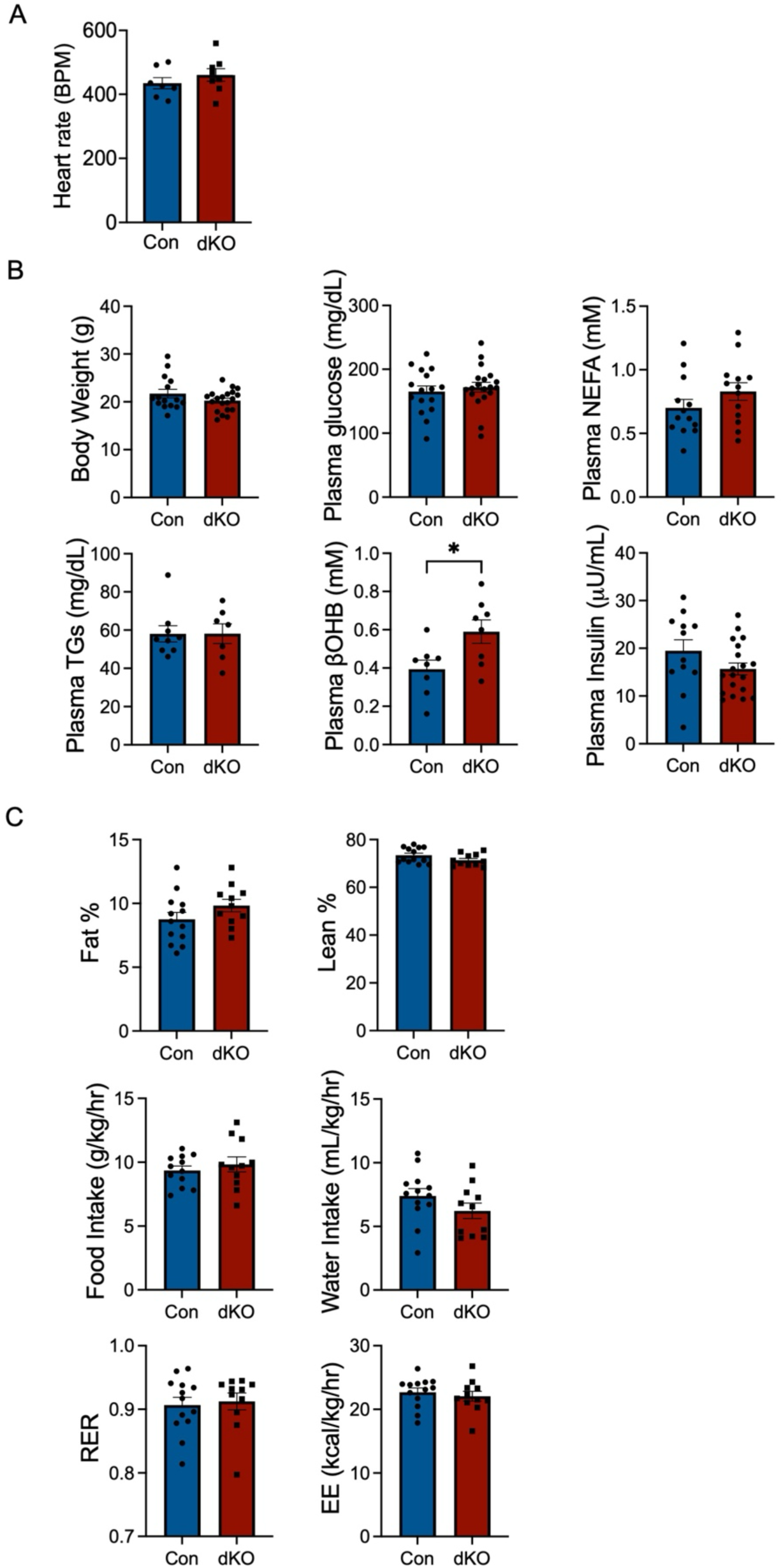
(A) Quantification of heart rate in beats per minute (BPM) from control and AMPK dKO mice at 6 weeks of age (n=7 per group). (B) Body weight and circulating plasma concentrations of glucose, non-esterified fatty acids (NEFA), triglycerides, β-hydroxybutyrate (βOHB), and insulin from control and atrial AMPK dKO mice after 8 hours of fasting (n =15-20 per group). (C) Whole-body composition, food and water intake, respiratory exchange ratio, and energy expenditure measured in metabolic cages (EE: energy expenditure; RER: respiratory exchange ratio). Data are shown as mean ± SEM; Significance determined by 2-tailed Student’s *t* test. *p < 0.05 AMPK dKO vs. control.

**Supplemental Figure 2.**
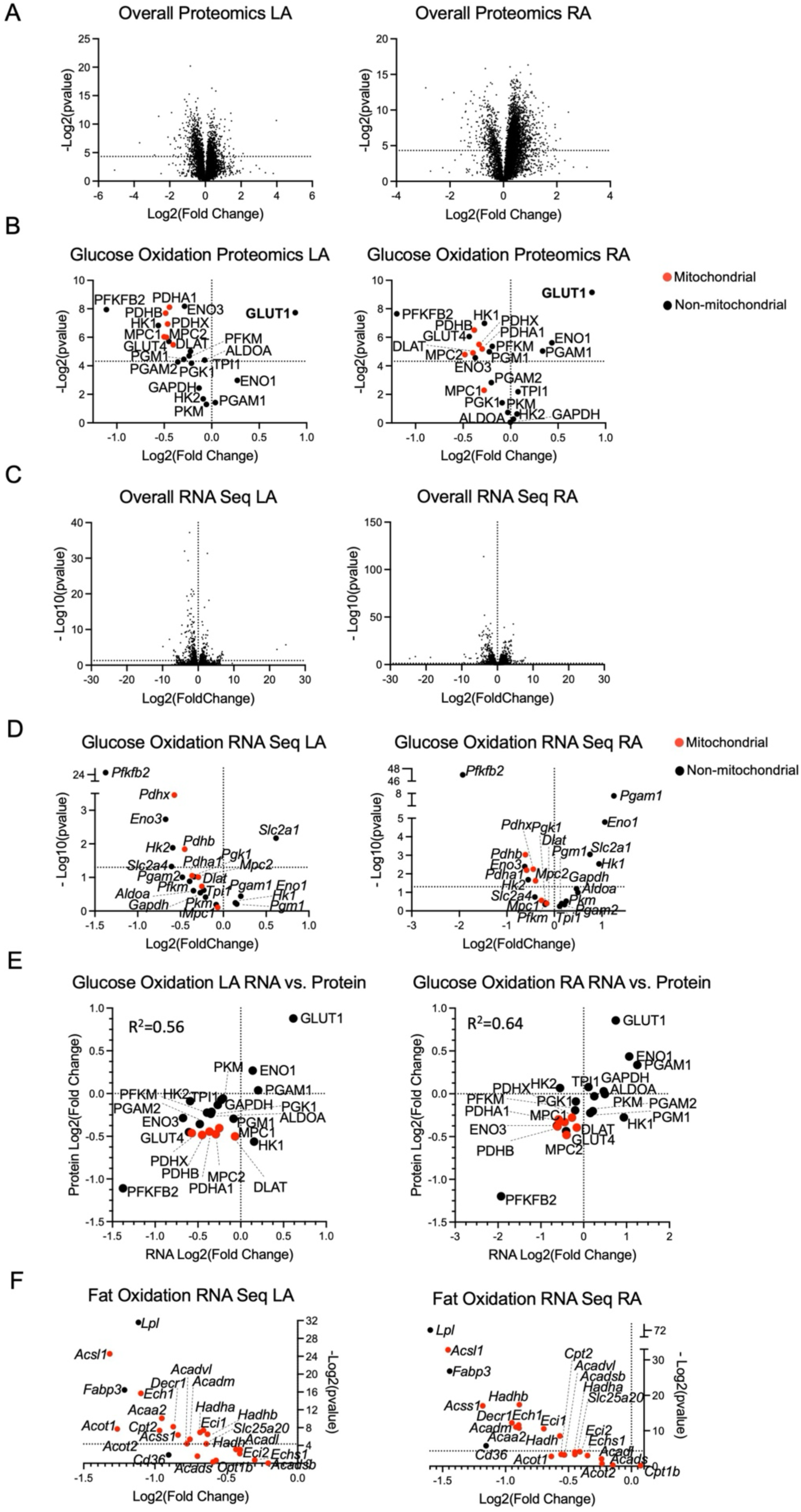
(A) Proteomic analysis of all proteins measured (6609 unique proteins), comparing atrial control and AMPK dKO mice at 6 weeks (n=3 per group), for left atria (LA) and right atria (RA). Data show fold-change (AMPK dKO vs. control) and statistical significance between groups. Graph includes dashed line that indicates p=0.05. (B) Proteomic analysis of glucose metabolism protein expression, comparing atrial control and AMPK dKO mice for LA and RA. Data show fold-change (AMPK dKO vs. control) and statistical significance between groups. Proteins localized to mitochondria are indicated in red, and other proteins in black. (C) RNA-sequencing (RNA-seq) analysis of all transcripts measured (52535 unique transcripts), comparing atrial control and AMPK dKO mice at 6 weeks (n=3 per group), for left and right atria. Data show fold-change (AMPK dKO vs. control) and statistical significance between groups. (D) Quantification of changes in transcripts for proteins involved in in glucose metabolic, comparing atrial control and AMPK dKO mice at 6 weeks, for LA and right atria RA. (E) Correlation between the degree of downregulation of RNA transcripts (RNA-seq) and protein expression (proteomic analysis) of components of the glucose metabolic pathway, comparing control and AMPK dKO mice, for LA and RA. (F) RNA-seq analysis of transcripts encoding proteins regulating fatty acid metabolism, comparing control and atrial AMPK dKO mice at 6 weeks, for left and right atria. Data show fold-change (AMPK dKO vs. control) and statistical significance between groups.

**Supplemental Figure 3.**
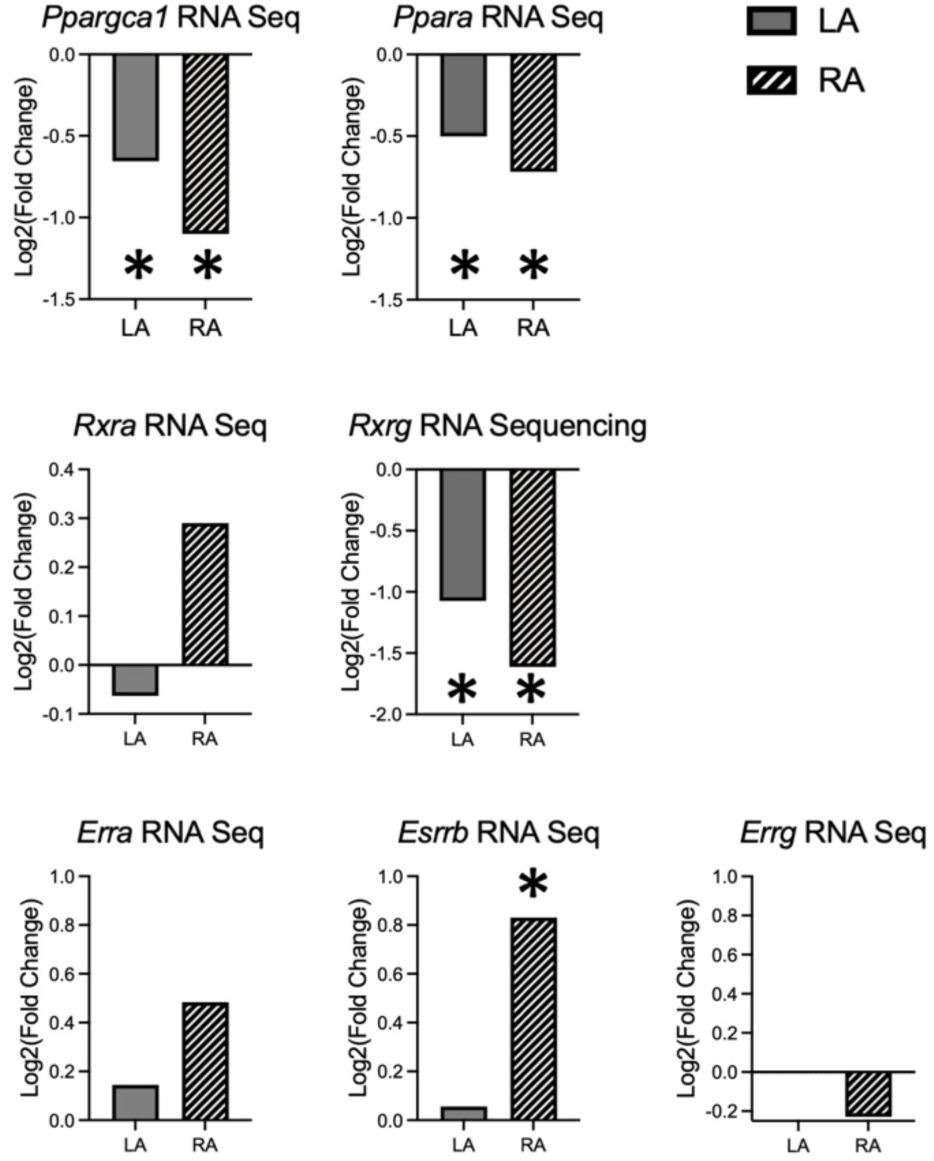
RNA-seq analysis comparing transcripts encoding transcription factors for proteins involved in fatty acid oxidation (*Ppargca1*, *Ppara, Rxra, Rxrg, Esrra, Esrrb,* and *Esrrg*) in control and atrial AMPK dKO mice. Data show fold-change (AMPK dKO vs. control) for left atria (LA) and right atria (RA) and indicate statistical significance between groups. Significance determined by 2-tailed Student’s *t* test. *p < 0.05 AMPK dKO vs. control.

